# The condensin complex is a mechanochemical motor that translocates along DNA

**DOI:** 10.1101/137711

**Authors:** Tsuyoshi Terekawa, Shveta Bisht, Jorine M. Eeftens, Cees Dekker, Christian H. Haering, Eric C. Greene

**Author notes:** Equal contribution (Tsuyoshi Terekawa, Shveta Bisht, Jorine M. Eeftens).

## Abstract

**One Sentence Summary:** Single-molecule imaging reveals that eukaryotic condensin is a highly processive DNA-translocating motor complex.

**Abstract:** Condensin plays crucial roles in chromosome organization and compaction, but the mechanistic basis for its functions remains obscure. Here, we use single-molecule imaging to demonstrate that *Saccharomyces cerevisiae* condensin is a molecular motor capable of ATP hydrolysis-dependent translocation along double-stranded DNA. Condensin’s translocation activity is rapid and highly processive, with individual complexes traveling an average distance of ≥10 kilobases at a velocity of ∼60 base pairs per second. Our results suggest that condensin may take steps comparable in length to its ∼50-nanometer coiled-coil subunits, suggestive of a translocation mechanism that is distinct from any reported DNA motor protein. The finding that condensin is a mechanochemical motor has important implications for understanding the mechanisms of chromosome organization and condensation.

---

Structural maintenance of chromosomes (SMC) complexes are the major organizers of chromosomes in all living organisms (1, 2). These protein complexes play essential roles in sister chromatid cohesion, chromosome condensation and segregation, DNA replication, DNA damage repair, and gene expression. A distinguishing feature of SMC complexes is their large ring-like architecture, the circumference of which is made up of two SMC protein coiled-coil proteins and a single kleisin subunit (Fig.1A) (1-4). The ∼50-nm long antiparallel coiled-coils are connected at one end by a stable dimerization interface, referred to as the hinge domain, and at the other end by globular ATP-binding cassette (ABC) family ATPase domains (5). The ATPase domains are bound by a protein of the kleisin family, along with additional accessory subunits, which vary for different types of SMC complexes (Fig. 1A). The relationship between SMC structures and their functions in chromosome organization is not completely understood (6), but many models envision that the coiled-coil domains allow the complexes to topologically embrace DNA (1-4). Given the general resemblance to myosin and kinesin, some early models postulated that SMC proteins might be mechanochemical motors (7-10).

**Fig. 1.**
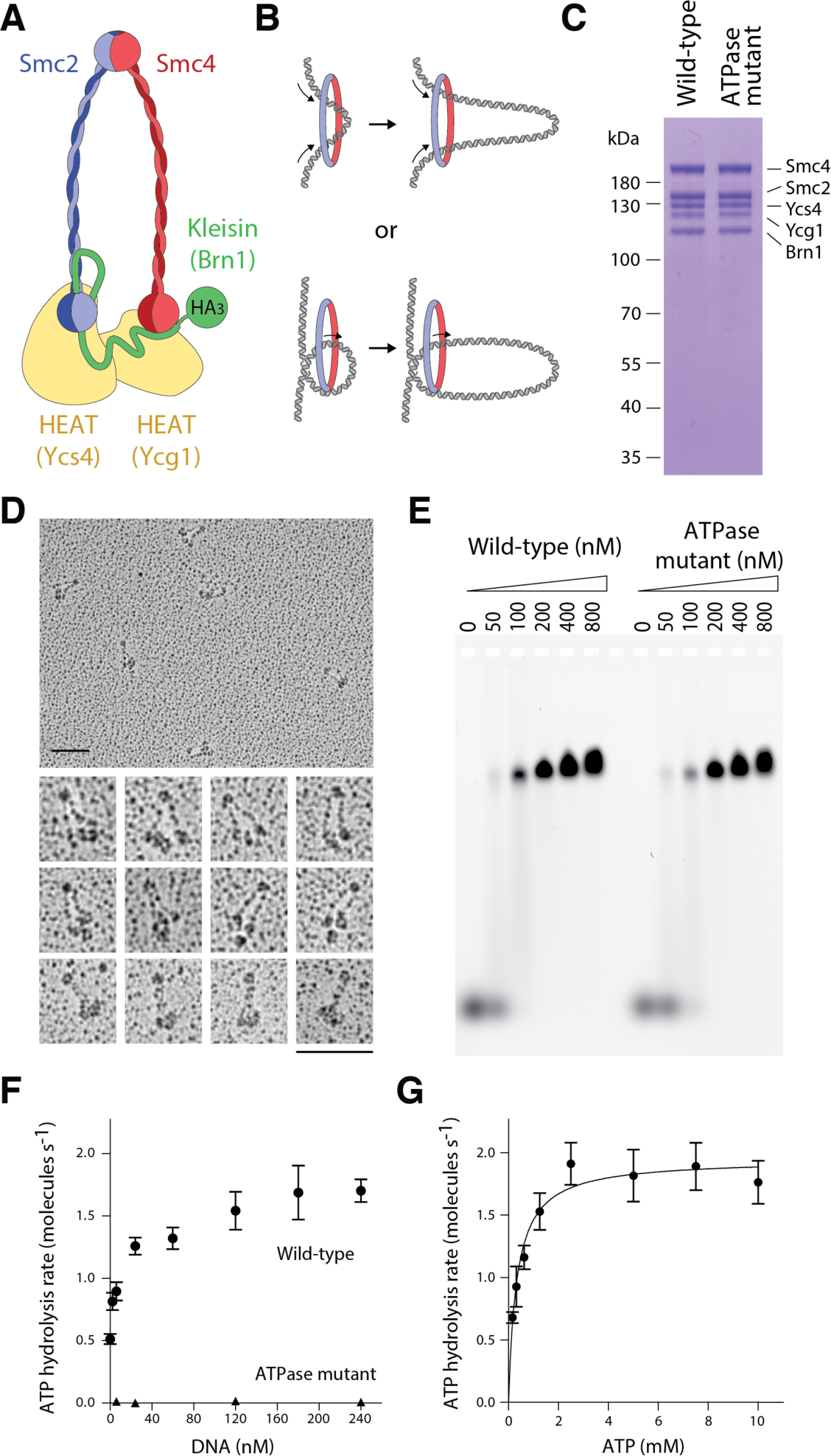
Biochemistry of budding yeast condensin holocomplexes. **A**) Schematic of the *S. cerevisiae* condensin complex. The Brn1 kleisin subunits connects the ATPase head domains of the Smc2–Smc4 heterodimer and recruits the HEAT-repeat subunits Ycs4 and Ycg1. The cartoon highlights the position of the HA_3_-tag used for labeling. (**B**) Conceptual schematic of loop extrusion models. (**C)** Wild-type and ATPase-deficient Smc2(Q147L)–Smc4(Q302L) condensin complexes analyzed by SDS PAGE and Coomassie staining. **(D)** Electron micrographs of wild-type condensin holocomplexes rotary shadowed with Pt/C. Scale bar: 100 nm. **(E)** Electrophoretic mobility shift assays with 6-FAM labelled 45-bp dsDNA substrate (100 nM) and the indicated protein concentrations. **(F)** ATP hydrolysis by wild-type and ATPase mutant condensin complexes (0.5 μM) upon addition of increasing concentrations of a 6.4-kb linear DNA at saturated ATP concentrations (5 mM). The plot shows mean ± standard deviation of *n* = 3 (wild-type) or 2 (ATPase mutant) independent experiments. **(G)** Michaelis-Menten kinetics for the rate of ATP hydrolysis by wild-type condensin complexes (0.5 μM) at increasing ATP concentrations in the presence of 240 nM 6.4-kb linear DNA. The plot shows mean ± S.D. of N = 3 independent experiments. The fit corresponds to a *K*_m_ of 0.4 ± 0.07 mM for ATP and *k*_*cat*_ of 2.0 ± 0.1 s^-1^ per molecule of condensin (mean ± standard error).

SMC complexes are thought to regulate genome architecture by physically linking distal chromosomal loci, but how these bridging interactions might be formed remains unknown (1, 2, 11). An early model suggested that many three-dimensional (3D) features of eukaryotic chromosomes might be explained by DNA loop extrusion (Fig. 1B) (12), and recent polymer dynamics simulations have shown that loop extrusion can recapitulate the formation of topologically associating domains (TADs), chromatin compaction, and sister chromatid segregation (13-17). This loop extrusion model assumes a central role for SMC complexes in actively creating the DNA loops (11, 12). Similarly, it has been proposed that prokaryotic SMC proteins may structure bacterial chromosomes through an active loop extrusion mechanism (18). Yet, the loop extrusion model remains hypothetical, in large part because the motor activity that is necessary for driving loop extrusion could not be identified (11). Indeed, the absence of an identifiable motor activity in SMC complexes instead has lent support to alternative models in which DNA loops are not actively extruded, but instead are captured and stabilized by stochastic pairwise SMC binding interactions to bridge distal loci (19).

To help distinguish between possible mechanisms of SMC protein-mediated chromosomal organization, we examined the DNA-binding properties of condensin (20). We overexpressed the five subunits of the condensin complex in budding yeast and purified the complex to homogeneity (Fig. 1C and fig. S1). Electron microscopy images confirmed that the complexes were monodisperse (Fig. 1D). As previously described for electron micrographs of immunopurified *Xenopus laevis* or human condensin (21), we observed electron density that presumably corresponds to the two HEAT-repeat subunits in close vicinity of the Smc2–Smc4 ATPase head domains. We confirmed that the *S. cerevisiae* condensin holocomplex binds double-stranded (ds) DNA and hydrolyzes ATP *in vitro* (Fig. 1E and F). Addition of dsDNA stimulated the condensin ATPase activity ∼3-fold, which is consistent with previous measurements with *X. laevis* condensin I complexes (22), and revealed *K*_*m*_ and *k*_*cat*_ values of 0.4 ± 0.07 mM and 2.0 ± 0.1 s^-1^, respectively, for ATP hydrolysis in the presence of linear dsDNA (Fig. 1G). An ATPase-deficient version of condensin with mutations in the γ-phosphate switch loops (Q-loops) of Smc2 and Smc4 still bound DNA (Fig. 1E), but exhibited no ATP hydrolysis activity (Fig. 1F).

We then used total internal reflection fluorescence microscopy (TIRFM) to visualize condensin binding to DNA curtains at the single-molecule level (23). Initial experiments using single-tethered DNA curtains demonstrated that condensin could promote extensive ATP hydrolysis-dependent DNA compaction, which was reversible by increasing the salt concentration to 0.5 M NaCl (Fig. S2). We next asked whether we could directly visualize the binding of single fluorescently-tagged condensin holocomplexes to double-tethered DNA substrates. We fluorescently labeled condensin with quantum dots (Qdots) conjugated to antibodies against the HA_3_-tag fused to the Brn1 kleisin subunit (Fig. 1A). Electrophoretic mobility shift assays confirmed that condensin was quantitatively labeled (Fig. S3A). Importantly, binding to the Qdots inhibited neither condensin’s ATP hydrolysis activity nor its ability to alter DNA topology (Fig. S3B and C). We prepared double-tethered curtains by attaching the DNA substrates (∼48.5-kb *λ*-DNA) to a supported lipid bilayer through a biotin-streptavidin linkage, and aligned one end of the DNA molecules at nanofabricated chromium (Cr) barriers and anchored the other end of the DNA to Cr pedestals located 12 μm downstream (Fig. 2A) (23).

**Fig. 2.**
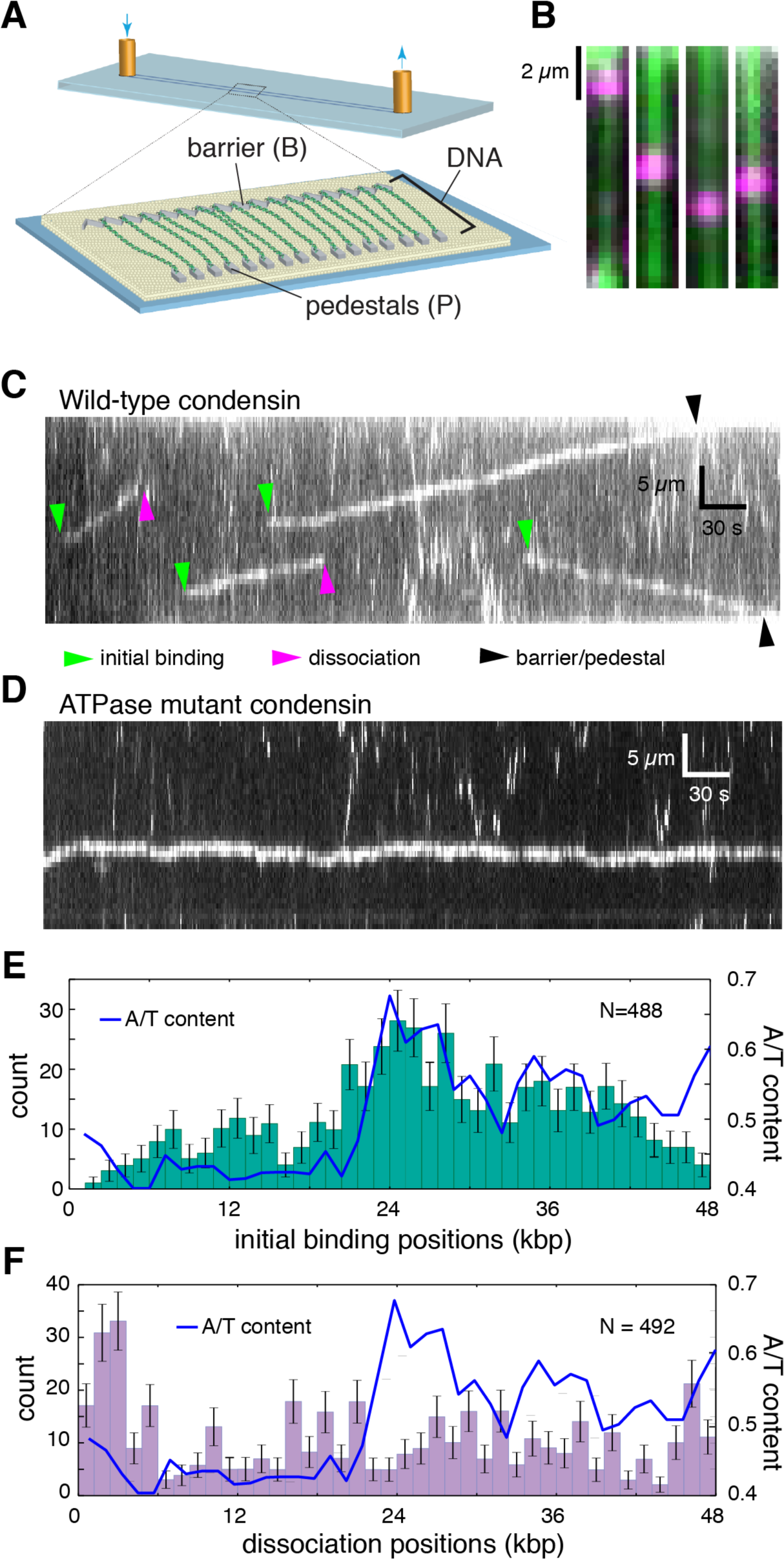
DNA curtain assay for condensin DNA-binding activity. (**A**) Schematic of the double-tethered DNA curtain assay. (**B**) Still images showing Qdot-tagged condensin (magenta) bound to YoYo1-stained DNA (green). (**C**) Kymograph showing examples of Qdot-tagged condensin translocating on a single DNA molecule (unlabeled); the initial condensin binding sites, dissociation positions, and collisions with the barriers/pedestals are highlighted with color-coded arrowheads. (**D**) Kymograph showing Qdot-tagged ATPase-deficient mutant Smc2(Q147L)– Smc4(Q302L) condensin undergoing 1D diffusion on DNA (unlabeled). (**E**) Initial binding sites and (**F**) dissociation site distributions of condensin superimposed on the AT content of the *λ*-DNA substrate. All reactions contained 4 mM ATP. Error bars in (**E**) and (**F**) represent standard deviations calculated by boot strap analysis.

Using double-tethered curtains, we were able to detect binding of condensin complexes to individual DNA molecules (Fig. 2B). Kymographs revealed that, remarkably, ∼85% of all bound condensin complexes (N = 671) underwent linear motion along the DNA (Fig. 2C). The up/down direction of movement was observed to be random, but once a complex started translocation, it did generally proceed unilaterally without a reversal of direction (reversals were observed occasionally, in 6% of the traces). Condensin has not been previously shown to act as a molecular motor, but the observed movement is fully consistent with expectations for an ATP-dependent DNA-translocating motor protein. The ATPase-deficient Q-loop mutant of condensin only exhibited motion consistent with random 1-dimensional (1D) diffusion (Fig. 2D). Wild-type condensin in the presence of the non-hydrolyzable ATP analog ATPγS similarly only displayed 1D-diffusion (Fig. S4A). Analysis of the initial binding positions for wild-type condensin revealed a preferential binding to A/T-rich regions (Pearson’s *r* = 0.66, *P* = 5×10^−6^; Fig. 2E), similar to values reported for *Schizosaccharomyces pombe* cohesin (24). In contrast, the condensin dissociation positions were not correlated with A/T content (Pearson’s *r* = –0.05, *P* = 0.77), nor were there any other preferred regions for dissociation within the DNA, with the exception of the Cr barriers and pedestals (Fig. 2F). These findings are consistent with a model where condensin loads at A/T-rich sequences and then translocates away. Interestingly, previous single-molecule experiments demonstrated rapid 1D diffusion of cohesin on DNA, but found no evidence for ATP-dependent translocation, suggesting that there may be a difference between how the two SMC complexes process DNA (24, 25).

We used particle tracking to quantitatively analyze the movement of condensin on DNA (Fig. 3A and B, and fig. 4SB). Wild-type condensin did not travel in a preferred direction: 52% (N=255/491) went one direction, and 48% went the opposite direction (N=236/491). The condensin ATPase mutant did not exhibit any evidence of unidirectional translocation. Mean squared displacement (MSD) plots generated from condensin tracking data revealed increasing slopes (Fig. 3C), which is only consistent with directed motion (26). In contrast, MSD plots were linear for the ATPase-deficient condensin mutant (Fig. 3D) and for wild-type condensin in the presence of ATPγS (Fig. S4C). Linear MSD plots were characteristic of random diffusive motion (26), yielding diffusion coefficients (*D*_1,*obs*_) of (1.7 ± 1.4)×10^−3^ and (0.8 ± 1.0)×10^−3^ μm^2^ s^-1^ (mean±S.D.) for ATPase-deficient condensin and wild-type condensin plus ATPγS, respectively.

**Fig. 4.**
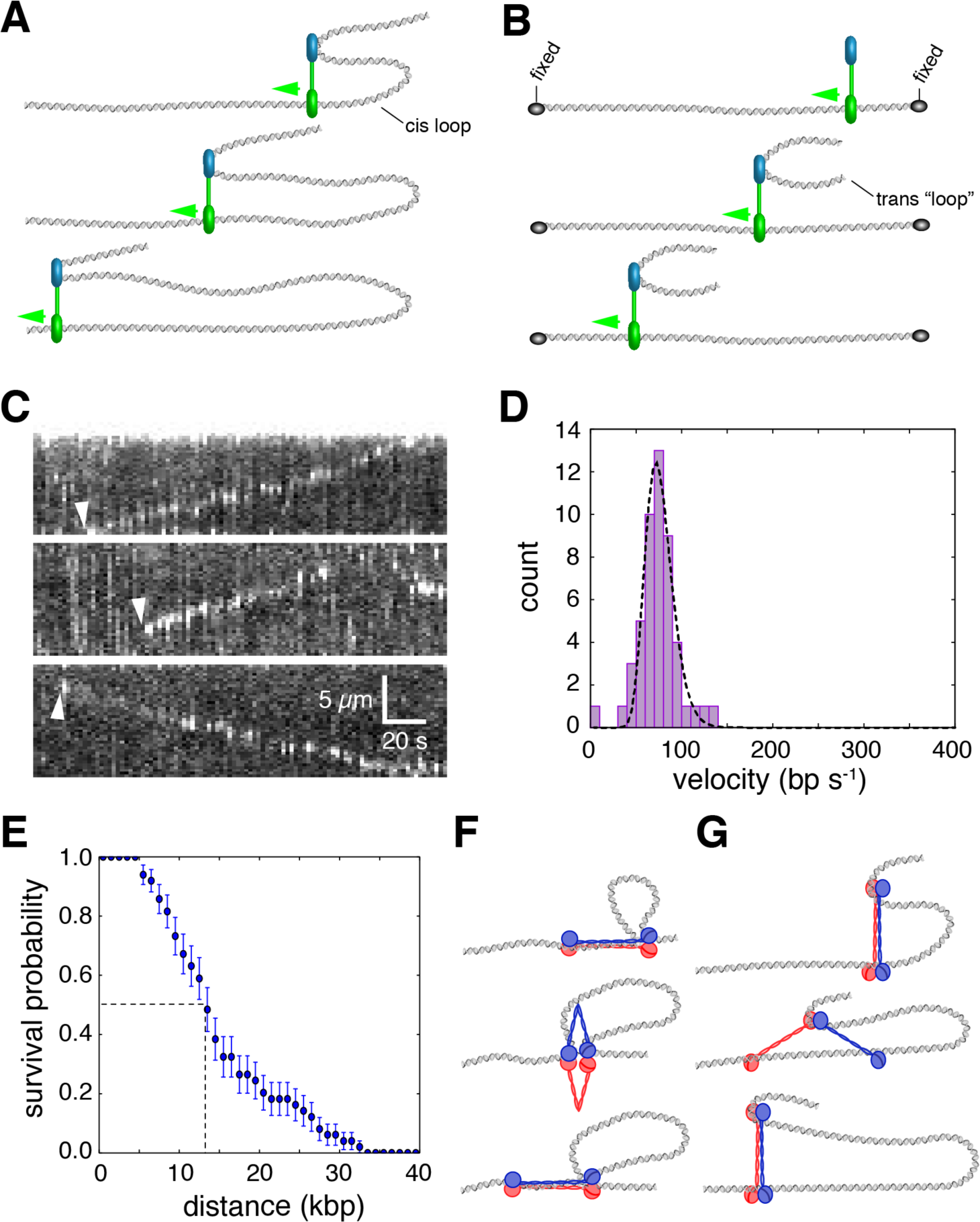
Coupling condensin motor activity to DNA loop extrusion. (**A**) Minimal mechanistic framework necessary for coupling ATP-dependent translocation to the extrusion of a *cis* DNA loop. In this generic model, a motor domain (green) must move away from a second DNA-binding domain (blue), and this secondary domain can either remain stationary (as depicted) or it may also act as a motor domain and move in the opposite direction (not depicted). (**B**) Detection of *Cis* loop extrusion is not possible when the DNA is held in a fixed configuration as in the double-tethered curtains that allows for direct detection of condensin motor activity in the absence of condensation (top panel). The middle and bottom panels show a schematic of an assay to mimic *cis* DNA loop extrusion by providing a second *λ*-DNA substrate in *trans*. (**C**) Examples of kymographs showing translocation of a second *λ*-DNA substrate (stained with YoYo1) provided in *trans* in the presence of unlabeled condensin. The presence of the *trans* DNA substrate is revealed as regions of locally high YoYo1 signal intensity, as highlighted by arrowheads. The regions of higher signal intensity are not detected when the trans DNA is omitted from the reaction. (**D**) Velocity distribution histogram and (**E**) survival probability plot for condensin bound to the *trans* DNA substrate. The dashed line in (**E**) highlights the translocation distance corresponding to dissociation of one half of the bound condesin complexes. Error bars represent standard deviations calculated by boot strap analysis. Cartoons of generalized models for condensin motor activity through a (**F**) “scrunching” or (**G**) “walking” mechanisms, both of which can be based upon ATP hydrolysis-dependent changes in the geometry of the SMC coiled-coil domains. Models are discussed in the text.

**Fig. 3.**
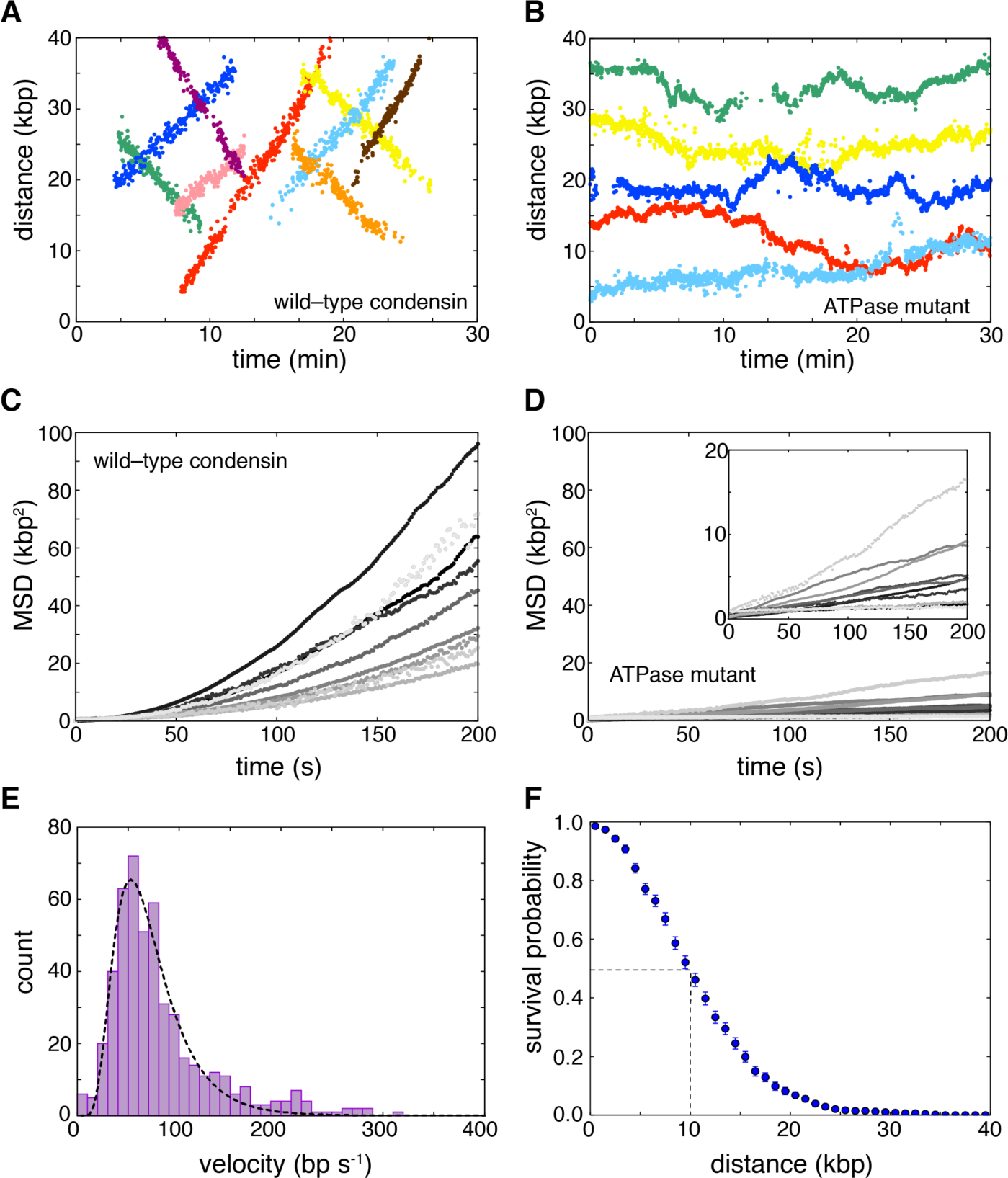
Condensin is an ATP-dependent mechanochemical molecular motor. **A**) Examples of tracked translocation trajectories for Qdot-tagged wild-type condensin and (**B**) the ATPase-deficient Smc2(Q147L)–Smc4(Q302L) condensin mutant. Mean squared displacement (MSD) plots for (**C**) wild-type condensin and (**D**) the ATPase deficient mutant obtained from the tracked trajectories. (**E**) Velocity distributions for condensin translocation activity; the dashed line is a log-normal fit to the translocation rate data. (**F**) Processivity measurements of condensin motor activity. The dashed line highlights the translocation distance corresponding to dissociation of one half of the bound condesin complexes. Error bars represent standard deviations calculated by boot strap analysis.

We used the tracking data to determine the velocity and processivity for wild-type condensin. A plot of the velocity distributions for data collected in the presence of saturated concentrations of ATP (4 mM; Fig. 1G) was well described by a log-normal distribution, revealing a mean apparent translocation velocity of 63 ± 36 bp s^-1^ (16 ± 9 nm s^-1^) (mean±S.D.) (Fig. 3E). Interestingly, upon initial binding, condensin paused for a brief period (τ_*pause*_=13.3 ± 1.5 s) before beginning to move along the DNA, which suggests the existence of a rate-limiting step prior to becoming active for translocation (Fig. 2C and fig. S5). Each translocating condensin complex remained bound to the DNA for an average total time of 4.7 ± 0.2 min and traveled on average 10.3 ± 0.4 kb (2.6 ± 0.1 μm) before dissociating (Figs. 3F and fig. S6A). This value provides merely a lower limit of the processivity of condensin, because a significant fraction (42%) of the complexes traveled all the way to the ends of the 48.5-kb *λ*-DNA, where they collided with the Cr barriers or pedestals (see, for example, Fig. 2C). There was no correlation between translocation velocity and processivity at a given ATP concentration (Pearson’s *r* = 0.035, *P* = 0.43 at 4 mM ATP) (Fig. S6B). However, velocity and processivity both varied with ATP concentrations. Michaelis-Menten analysis revealed a *v*_*max*_ of 62 ± 2 bp s^-1^ and a *K*_*m*_ of 0.2 ± 0.04 mM ATP (Fig. S7A and B). The initial pause time (τ_*pause*_) also varied with ATP concentration, from a mean value of 3.9 ± 0.8 min at 50 μM ATP to 13.3 ± 1.5 s at 4 mM ATP, suggesting that this delay reflects a transition from a translocation-inactive to translocation-active state that is dependent upon ATP binding, ATP hydrolysis, or both (Fig. S7C).

Our finding that condensin is an ATP hydrolysis-dependent molecular motor lends support to models invoking SMC protein-mediated loop extrusion as a means for 3D genome organization. An important prediction of the loop extrusion model is that condensin must simultaneously interact with two distal regions of the same chromosome, and at least one (or possibly both) of the interaction sites must translocate away from the other site, allowing for movement of the two contact points relative to one another (Fig. 4A) (12-16). The *“cis”* loop geometry is inaccessible in our double-tethered assays because the DNA is held in an extended configuration (Fig. 2B), which likely decouples loop extrusion from translocation. However, a *cis* loop configuration can be mimicked experimentally by providing a second DNA molecule in *trans* (Fig. 4B). To test the possible relationship between the observed linear translocation of condensin along the double-tethered DNA and the loop extrusion model, we asked whether condensin could move a second DNA substrate provided in *trans* relative to the tethered DNA. Indeed, fluorescently labeled (not extended) *λ*-DNA molecules added in *trans* moved at an apparent velocity of 66 ± 15 bp s^-1^ (16 ± 4 nm s^-1^, N=50) (Fig. 4C and D) while traveling an average distance of 14 ± 0.8 kb (3.4 ±0.3 μm, N=50) (Fig. 4E) – numbers that match excellently with the condensin motor properties measured above. These experiments strongly indicate that translocating condensin complexes were able to interact simultaneously with the tethered DNA and a second DNA, that condensin could translocate while bound to both DNA substrates, and that one piece of DNA was moved with respect to the other piece of DNA. We conclude that condensin is capable of moving two DNA substrates relative to one another, fulfilling a key expectation of the loop extrusion model.

Heretofore, a common argument against SMC proteins acting as molecular motors was their low rates of ATP hydrolysis when compared to other known nucleic acid motor proteins, which implied that they would not move fast enough to function efficiently on biologically relevant time scales. However, this discrepancy can be readily reconciled if condensin were able to take large steps, which is conceptually possible given its large 50-nm size. Based on the avaible data, it is in fact quite reasonable to invoke a large step size: Comparison of the single molecule translocation rate (∼60 bp s^-1^, or ∼14.9 nm s^-1^) to the bulk rate of ATP hydrolysis (*K*_*cat*_ = 2.0 s^-1^ in the presence of linear DNA) suggest that condensin may take steps on the order of ∼30 bp per molecule of ATP hydrolyzed. Even larger steps are inferred if each step is coupled to the hydrolysis of more than one molecule of ATP. These estimates assume that all of the proteins are ATPase active (one would deduce a smaller step size if a fraction of the protein were inactive), and also assume perfect coupling between ATP hydrolysis and translocation (while a more inefficient coupling would necessitate even larger step sizes). The idea that condensin takes very large steps is also consistent with the step sizes reported from magnetic tweezer experiments of DNA compaction induced by *X. leavis* condensin (80 ± 40 nm, *Ref.* (27) or *S. cerevisiae* condensin (*J. Eeftens et al., unpublished*). Such large step sizes would seem to rule out models for condensin movement similar to common DNA motor proteins such as helicases, translocases or polymerases, which are typically found to move in 1-bp increments (28-31). Higher-resolution measurements may prove informative for further defining the fundamental step size for translocating condensin.

To explain our novel results, we search for possible models for condensin motor activity that (*i*) can explain the relationship between a slow ATP hydrolysis rate relative to the rate of translocation; (*ii*) can accommodate a very large step size; and (*iii*) are consistent with the physical dimensions of the SMC complex. Based on these criteria, we can think of two theoretical possibilities, both of which use the SMC coiled-coils as the means of motility: Condensin might translocate along DNA through reiterative extension and retraction of the long Smc2–Smc4 coiled-coil domains, allowing for movement through a “scrunching” mechanism involving rod- to butterfly-like structural transitions (Fig. 4F); or condensin may perhaps use a myosin- or kinesin-like “walking” mechanism (Fig. 4G). The maximum single step size for each model is defined by the physical dimensions of the SMC coiled-coils, corresponding to ≲50 nm and ≲100 nm for the scrunching and walking mechanisms, respectively (Fig. 4F and G). Both models are consistent with the range of condensin architectures observed by electron microscopy and atomic force microscopy (21, 32). Movements might be powered by similar ATPase-dependent transitions between different structural states as reported for prokaryotic SMC complexes (33-35), though it remains to be determined how conformational changes could be translated into the directed movement depicted in our models. Further refinement of the translocation mechanism will depend upon fully defining the structural transitions that take place during the ATP hydrolysis cycle and establishing a better understanding of whether and if so, how different domains in the condensin complex engage DNA.

The finding that condensin is a mechanochemical motor capable of translocating along DNA has important implications for understanding fundamental mechanisms of chromosome organization across all domains of life. We propose that the ATP hydrolysis-dependent motor activity of condensin may be intimately linked to its role in promoting chromosome condensation, suggesting that condensin, and perhaps other SMC proteins, may provide the driving forces necessary to support 3D chromosome organization and compaction through a loop extrusion mechanism. These findings raise the question of whether other types of SMC complexes also exhibit intrinsic motor activity, and what molecular or regulatory features distinguish SMC motor proteins from those SMC complexes that seemingly lack motor activity.

## Acknowledgements

We thank Damien D’Amours (Univ. of Montreal) for plasmids and yeast strains for overexpression of the condensin holocomplex, and members of the Haering, Greene, and Dekker laboratories for comments on the manuscript and the EMBL Electron Microscopy Facility and Proteomics Core Facilities for support. This work was funded by a MIRA grant from the National Institutes of Health to E.C.G. (R35GM118026), by EMBL and the ERC Consolidator grant to C.H.H. (ERC-2015-CoG 681365), and the ERC Advanced Grant SynDiv (ERC-ADG-2014 669598) and the Netherlands Organization for Scientific Research (NWO/OCW) (as part of the Frontiers of Nanoscience program) to C.D. T.T. was supported by a Japan Society for the Promotion of Science fellowship, and by a Uehara Memorial Foundation fellowship. S.B. was supported by an EMBL Interdisciplinary Postdoctoral fellowship (EIPOD) under Marie Curie Actions (COFUND). J.E. was supported by an EMBO short term fellowship.

## Author contributions

T.T. designed and conducted single-molecule experiments and data analysis. S.B. purified condensin complexes, and conducted bulk biochemical measurements and electron microscopy analysis. J.E. designed and implemented single-molecule experiments. All authors discussed the experimental findings and co-wrote the manuscript.

## Supplementary Materials

### Materials and Methods

***Condensin holocomplex overexpression and purification.*** The five subunits of the condensin complex were co-overexpressed in *Saccharomyces cerevisiae* from galactose-inducible promoters on 2μ high-copy plasmids (*URA3 leu2-d pGAL7-SMC4(wild-type or Q302L)-StrepII*_*3*_ *pGAL10-SMC2(wild-type or Q147L) pGAL1-BRN1-HA*_*3*_*-His*_*12*_ and *TRP1 leu2-d pGAL10-YCS4 pGAL1-YCG1*; yeast strains C4491 and C4724) as described (36), with the following modifications. Cultures were grown at 30°C in –URA-TRP dropout media containing 2% raffinose to OD_600_ of 1. Expression was induced with 2% galactose for 8 hours. Since expression of the Q-loop mutant complex affected the growth rate of the cultures, cells were initially grown at 30°C URA TRP dropout media containing glucose to OD_600_ of 1, transferred to media containing 2% raffinose for one hour and then induced by addition of galactose to 2%.

Cells were harvested by centrifugation, resuspended in buffer A (50 mM TRIS-HCl pH 7.5, 200 mM NaCl, 5% (v/v) glycerol, 5 mM β-mercaptoethanol, and 20 mM imidazole) containing 1× cOmplete EDTA-free protease-inhibitor mix (Roche) and lysed in a FreezerMill (Spex). The lysate was cleared by two rounds of 20 min centrifugation at 45,000 ×g at 4°C and loaded onto a 5 ml HisTrap column (GE Healthcare) pre-equilibrated with buffer A. The resin was washed with five column volumes buffer A containing 500 mM NaCl; buffer A containing 1 mM ATP, 10 mM KCl and 1 mM MgCl_2_; and then buffer A containing 40 mM imidazole to remove non-specifically bound proteins. Protein was eluted in buffer A containing 200 mM imidazole and transferred to buffer B (50 mM TRIS-HCl pH 7.5, 200 mM NaCl, 5% (v/v) glycerol, 1 mM DTT) using a desalting column. After addition of EDTA to 1 mM, PMSF to 0.2 mM and Tween20 to 0.01%, the protein was incubated with 2 ml (bed volume) of pre-equilibrated Strep-Tactin high-capacity Superflow resin (IBA).

The Strep-Tactin resin was packed into a column and washed with 15 resin volumes buffer B by gravity flow. Protein was eluted with buffer B containing 5 mM desthiobiotin. The eluate was concentrated by ultrafiltration and loaded onto a Superose 6 size exclusion chromatography column (GE Healthcare) pre-equilibrated in buffer B containing 1 mM MgCl_2_. Peak fractions were pooled and concentrated to 4 μM by ultrafiltration. Purified proteins were analyzed by SDS PAGE (NuPAGE 4-12% Bis-Tris protein gels, ThermoFisher Scientific) and protein bands were identified by in-gel digestion and mass spectrometric analysis.

#### Nick ligation assays

Nick ligation assays were performed as described earlier (37) with the following modifications. A 6.4-kb plasmid containing a single *BbvCI* nicking site was used as a substrate and relaxed by incubation with Nb.BbvCI (NEB). The nicking enzyme was heat-inactivated once the reaction was complete (as confirmed by agarose gel electrophoresis). For assessing supercoiling, reactions were set up in a volume of 20 μl containing 1 nM nicked plasmid DNA and varying amounts of condensin (7.8-500 nM) in 50 mM TRIS-HCl pH 7.5, 100 mM NaCl, 2.5% (v/v) glycerol, 10 mM MgCl_2_, 1 mM ATP and 10 mM DTT. Reactions were incubated at room temperature for 30 min before addition of 0.2 μl T4 ligase (5 Weiss U/μl, ThermoFisher Scientific) and fresh 1 mM ATP, followed by an additional 30 min incubation at room temperature to allow the ligation reaction to complete. Reactions were quenched by the addition of 60 μl stop buffer (50 mM TRIS-HCl pH 7.5, 10 mM EDTA, 1% SDS, 100 μg/ml proteinase K) and incubation at 37°C for 30 min. DNA was purified by phenol-chloroform extraction and ethanol precipitation. The DNA pellet was re-suspended in TE buffer and topoisomers were resolved at 4 V/cm for 9 h on a 0.7% TAE agarose gel containing 0.2 μg/ml chloroquine. The gel running buffer was also supplemented with chloroquine at the same concentration. Gels were stained with ethidium bromide and scanned on a Typhoon FLA 9500 (GE Healthcare).

#### Pt/C rotary shadowing electron microscopy

Samples for platinum/carbon (Pt/C) shadowing were prepared following the glycerol spray method (38). Condensin samples were diluted to a concentration of 0.05 μM in freshly prepared 200 mM NH_4_HCO_3_ pH 7.5, 30% (v/v) glycerol and 1 mM DTT, immediately sprayed onto freshly cleaved mica and dried under vacuum. Pt/C was shadowed at an angle of 7° followed by deposition of a stabilizing layer of carbon. The Pt/C layers were then floated off and placed onto 100 mesh copper grids. The grids were dried and imaged on a Morgagni TEM (FEI).

#### Electrophoretic mobility assay

6-carboxyfluorescein (6-FAM) labelled 45 bp dsDNA was prepared by annealing two complementary HPLC-purified DNA oligonucleotides (IDT, 5’-6-FAM-CCA GCT CCA ATT CGC CCT ATA GTG AGT CGT ATT ACA ATT CAC TGG-3’; 5’- CCA GTG AAT TGT AAT ACG ACT CAC TAT AGG GCG AAT TGG AGC TGG-3’) in annealing buffer (10 mM TRIS-HCl pH 7.5, 50 mM NaCl, 1 mM EDTA) at a concentration of 10 μM in a temperature gradient of 0.1°C/s from 95°C to 4°C. 10 μl reactions were prepared with 100 nM 6-FAM-dsDNA and protein concentrations ranging from 50 to 800 nM in reaction buffer (40 mM TRIS–HCl pH 7.5, 125 mM NaCl, 5 mM MgCl_2_, 10 % glycerol, and 1 mM DTT). After 10 min incubation at room temperature (∼25°C), free DNA and DNA–protein complexes were resolved by electrophoresis for 10 h at 2 V/cm on 0.8% (w/v) TAE-agarose gels at 4°C. 6-FAM labelled DNA was detected on a Typhoon FLA 9500 scanner (GE Healthcare) with excitation at 473 nm and detection using a 510-nm long pass filter.

#### Native gel electrophoresis

Protein samples (100 nM) were incubated with 1X and 2X molar ratio of anti-HA Qdots for 10 min at room temperature and the loaded onto a composite agarose-acrylamide gel (0.5% agarose and 2% acrylamide) (39). Electrophoresis was performed in TBE buffer at 30V for 10 h at 4°C. The gels were analyzed by silver staining.

#### ATP hydrolysis assays

ATPase reactions were set up in a volume of 5 μl containing 0.1 μM or 0.5 μM condensin and the indicated concentrations of DNA in 40 mM TRIS-HCl pH 7.5, 125 mM NaCl, 5 mM MgCl_2_, 0.5 mg/ml BSA and 1 mM DTT. 6.4-kb plasmid DNA had been linearized by *NheI* restriction digest and purified by phenol-chloroform extraction and ethanol precipitation. Reactions were initiated by the addition of ATP at the indicated concentrations (containing 6.7 nM [α-^32^P]ATP) and incubated at room temperature. At consecutive time intervals, 1 μl of the reaction mix was spotted onto PEI cellulose F TLC plates (Merck). TLC plates were developed in 0.5 M LiCl, and 1 M formic acid, exposed to imaging plates and analyzed on a Typhoon FLA 9500 scanner (GE Healthcare). ATP hydrolysis rates were calculated from the change of ATP/ADP ratios between time points in the linear range of the reaction.

#### Single molecule assays

Double-tethered DNA curtains were prepared as described previously (23, 24). Unless otherwise stated, all single molecule measurements were performed in condensin buffer (40 mM TRIS-HCl pH 7.5, 125 mM NaCl, 10 mM MgCl_2_, 1 mM DTT, 0.5 mg/mL BSA, and 4 mM ATP) and all assays were conducted at room temperature (∼25°C). Quantum dots were labeled with anti-HA antibodies as per the manufacturer’s instructions using a SiteClick^™^ Qdot*®* 705 Antibody Labeling Kit (ThermoFisher, Cat No. S10469). Purified condensin (1 μl of 1 μM stock) was labeled by mixing with 2 μl anti-HA quantum dots (1 μM) in 7 μl of condensin buffer and incubated on ice for 10 minutes. The labeling reactions were diluted to 100 μl with condensin buffer and then injected into the sample chambers at a flow rate 0.1 ml/min. Flow was then terminated, and the samples were incubated for an additional 20 minutes. Samples were visualized with a custom modified inverted Nikon microscope equipped with a Nikon 60× CFI Plan Apo VC water immersion objective, as described (23, 24). Image acquisition was initiated immediately before injecting condensin and continued throughout the 20-minute incubation. All images were acquired with an iXon EMCCD camera (Andor) at a 1 Hz frame acquisition rate. Note that in the absence of nucleotide co-factor, condensin adhered non-specifically to the surfaces of the sample chambers, so all single molecule measurements contained either ATP or ATPγS, as specified.

#### Particle tracking

The positions (*z*(*t*)) of each condensin complex were tracked using an in-house Python script. In this script, the intensity profile along DNA was fit with a one-dimensional Gaussian function, taking the mean of the Gaussian fits as the position (*z*(*t*)) in sub-pixel resolution (40). The total length of the *λ*-DNA substrate used in these experiments is 48,502 base pairs, or 16.49 μm. The DNA is extended to a mean length of ∼12 μm in the double-tethered DNA curtains, corresponding to ∼72% mean extension, and spans a distance of 48 pixels at 60× magnification. Were indicated, the measured length pixels was converted to base pairs by assuming that each pixel contains 1,010 base pairs of DNA, corresponding to a conversion factor of 4.04 base pairs per nanometer. All particle tracking data are measured in nanometers, and then converted to base pairs for comparison, and both sets of distances are reported. The mean square displacement (MSD) of each trajectory was calculated as *MSD* (Δ*t*) = 〈*zt*+ Δ*t* - *z*(*t*)〉. Translocation of each condensin complex showed a characteristic linear relationship between time and position (Fig. 3A). Thus, data points were fitted with a linear function to calculate slope, which corresponds to the translocation velocity. The translocation start (*t*_*s*_) and end (*t*_*e*_) times were manually obtained by visual inspection of the data, where the starting time was taken after the brief ∼13 s pausing time at the start of the linear trace. A small fraction (6%) of the condensin trajectories displayed a sudden change in direction, and in these instances the translocation end time (*t*_*e*_) was specified as the time when the molecules changed direction. The distance of translocation was defined as |*zt*_*e*_ − *z*(*t*_*s*_)|, and these values were used to calculate the survival probability plot (*i.e.* processivity) presented in Fig. 3F. The reported processivity values reflect the translocation distance at which one half of the condensin complexes dissociate from the DNA based upon the survival probability plots.

#### Single molecule trans loop assays

Assays were conducted using double-tethered DNA curtains, as described above. A 100 μL reaction mix was prepared in condensin buffer containing 1 nM condensin, 18 pM free *λ*-DNA (untagged), and 20 nM YoYo1 (ThermoFisher, Cat. No. Y3601). This reaction mix was then injected at a flow rate of 0.1 ml min^-1^ into a sample chamber that already contained double-tethered *λ*-DNA molecules. Note that the tethered *λ*-DNA was labeled at one end with biotin and at the other end with digoxigenin, as previously described (23, 24), whereas the free *λ*-DNA was not labeled. Buffer flow was then terminated, and the reactions were incubated for an additional 20 minutes at room temperature while capturing 100-millisecond images at 0.2 Hz frame acquisition rate. The laser was shuttered between each 100-millisecond exposure to minimize YoYo1-induced photo-damage. The resulting data were analyzed by particle tracking as describe above.

**Fig. S1.**
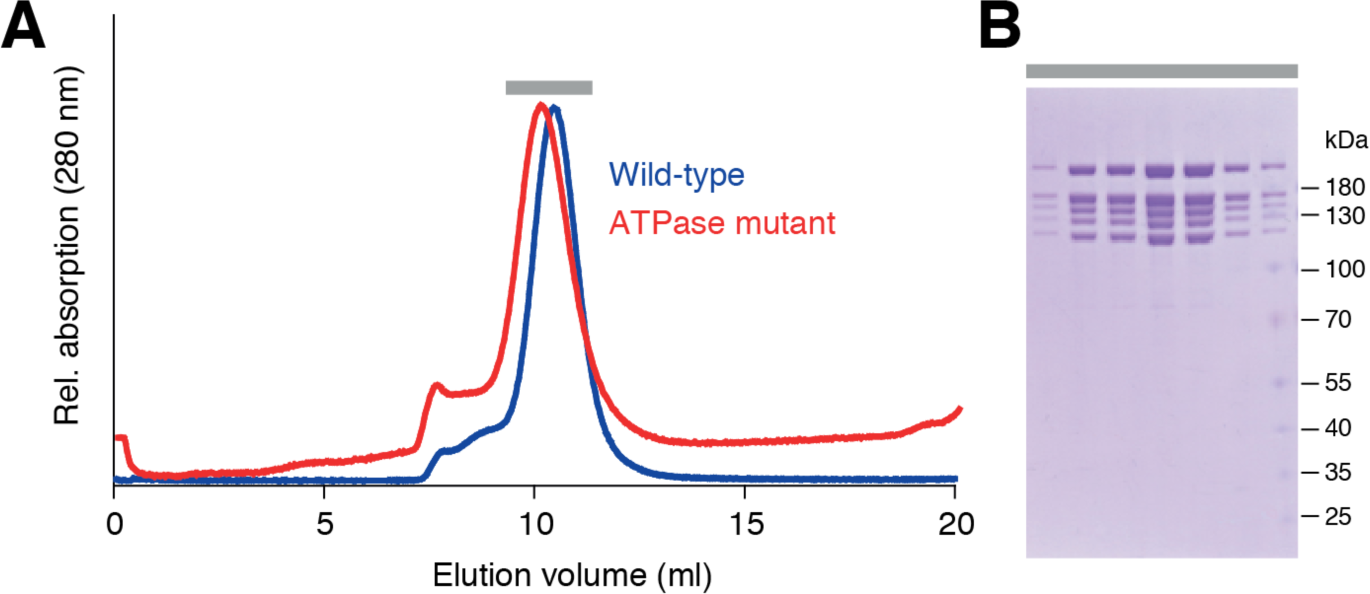
Purification of budding yeast condensin holocomplexes. **(A)** Size exclusion chromatograms of wild-type and ATPase-deficient Smc2(Q147L)–Smc4(Q302L) condensin complexes. (**B)** Analysis of peak fractions (grey bar) of the wild-type condensin purification by SDS PAGE and Coomassie staining.

**Fig. S2.**
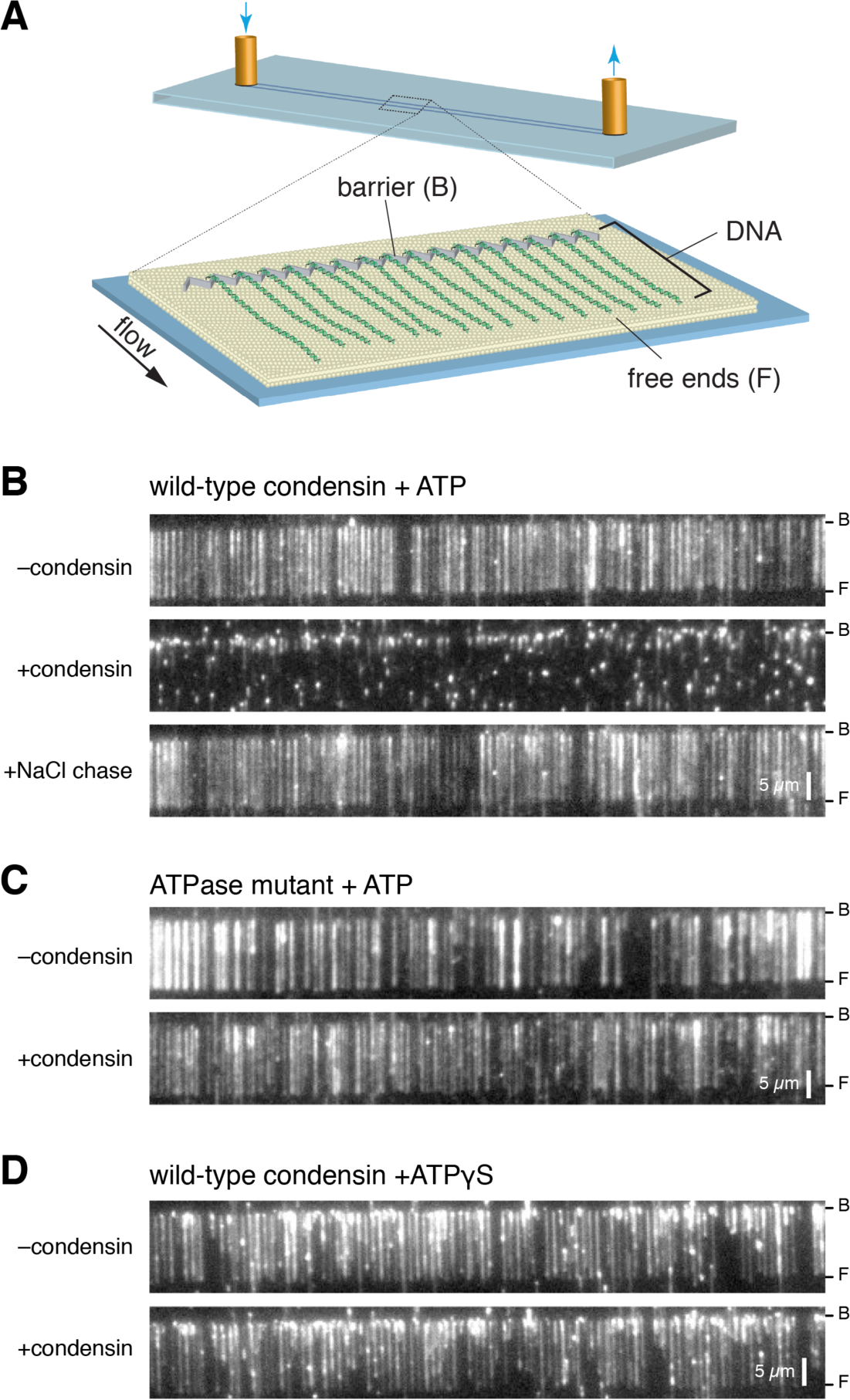
Condensin can reversibly compact DNA curtains. (**A**) Schematic of the single-tethered DNA curtain assay used to test for DNA compaction by unlabeled condensin (unlabeled). (**B**) Still images showing the YoYo1-stained DNA before addition of wild-type condensin, after a 20-minute incubation with 10 nM condensin and 4 mM ATP, and still images after chasing the reactions with 500 mM NaCl. (**C**) Still images showing the YoYo1-stained DNA before addition of ATPase deficient condensin, after a 20-minute incubation (in the absence of buffer flow) with 10 nM ATPase-deficient condensin mutant and 4 mM ATP. (**D**) Still images showing the YoYo1-stained DNA before addition of wild-type condensin, after a 20-minute incubation with 10 nM condensin and 4 mM ATPγS.

**Fig. S3.**
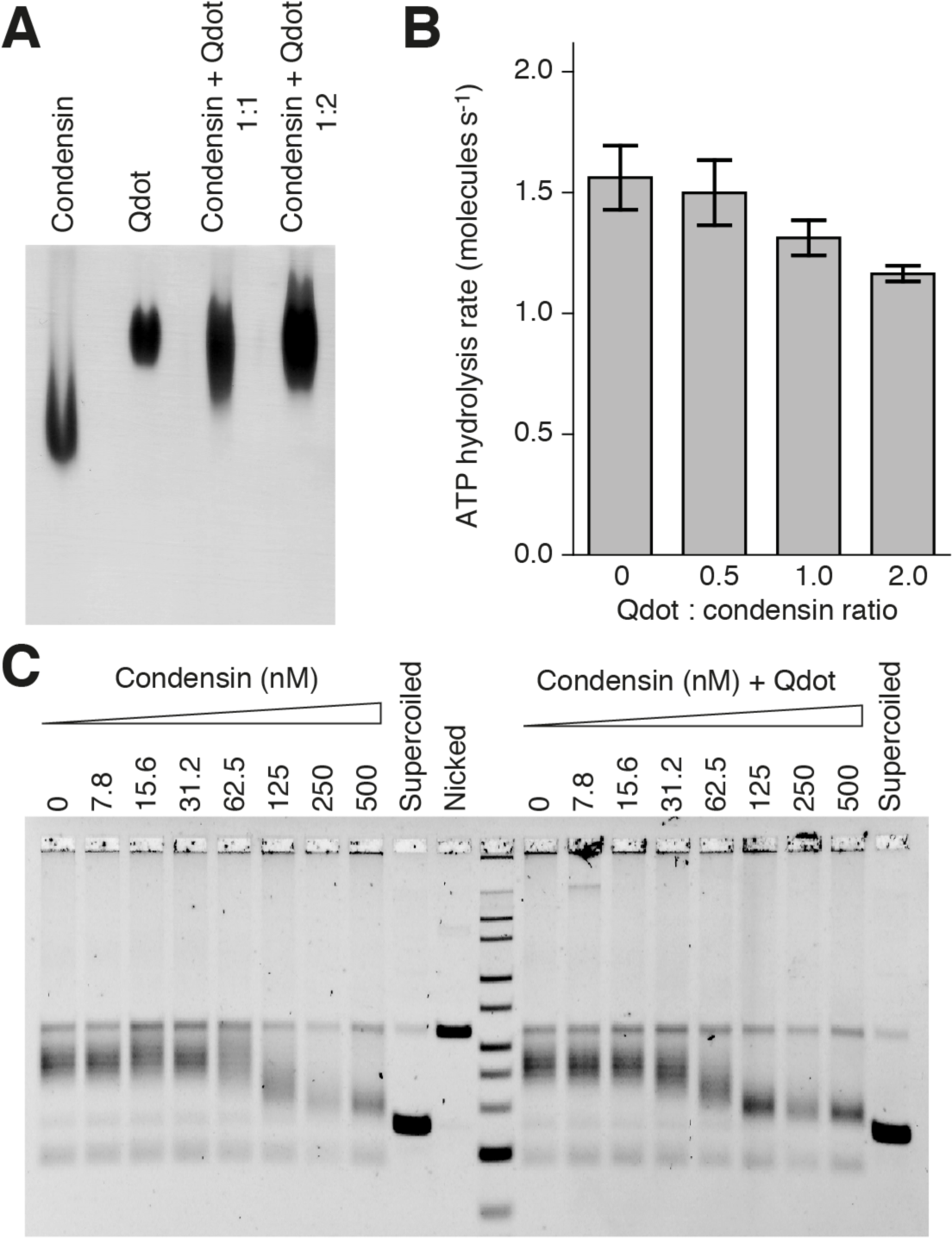
Condensin labeling with quantum dots. **(A)** Native composite agarose-acrylamide gel electrophoresis of wild-type condensin complexes upon addition of Qdots coupled to antibodies directed against the HA_3_ epitope tag at the C terminus of the Brn1 kleisin condensin subunit. (**B)** Effect of increasing ratios of anti-HA Qdot on the ATPase hydrolysis rate by wild-type condensin complexes (0.1 μM) in the presence of 6.4-kb linear DNA (240 nM) at saturated ATP concentrations (5 mM). (**C)** Nick ligation assay of a 6.4-kb circular DNA (1 nM) with wild-type condensin complexes alone and in the presence of an equimolar amount of anti-HA Qdot.

**Fig. S4.**
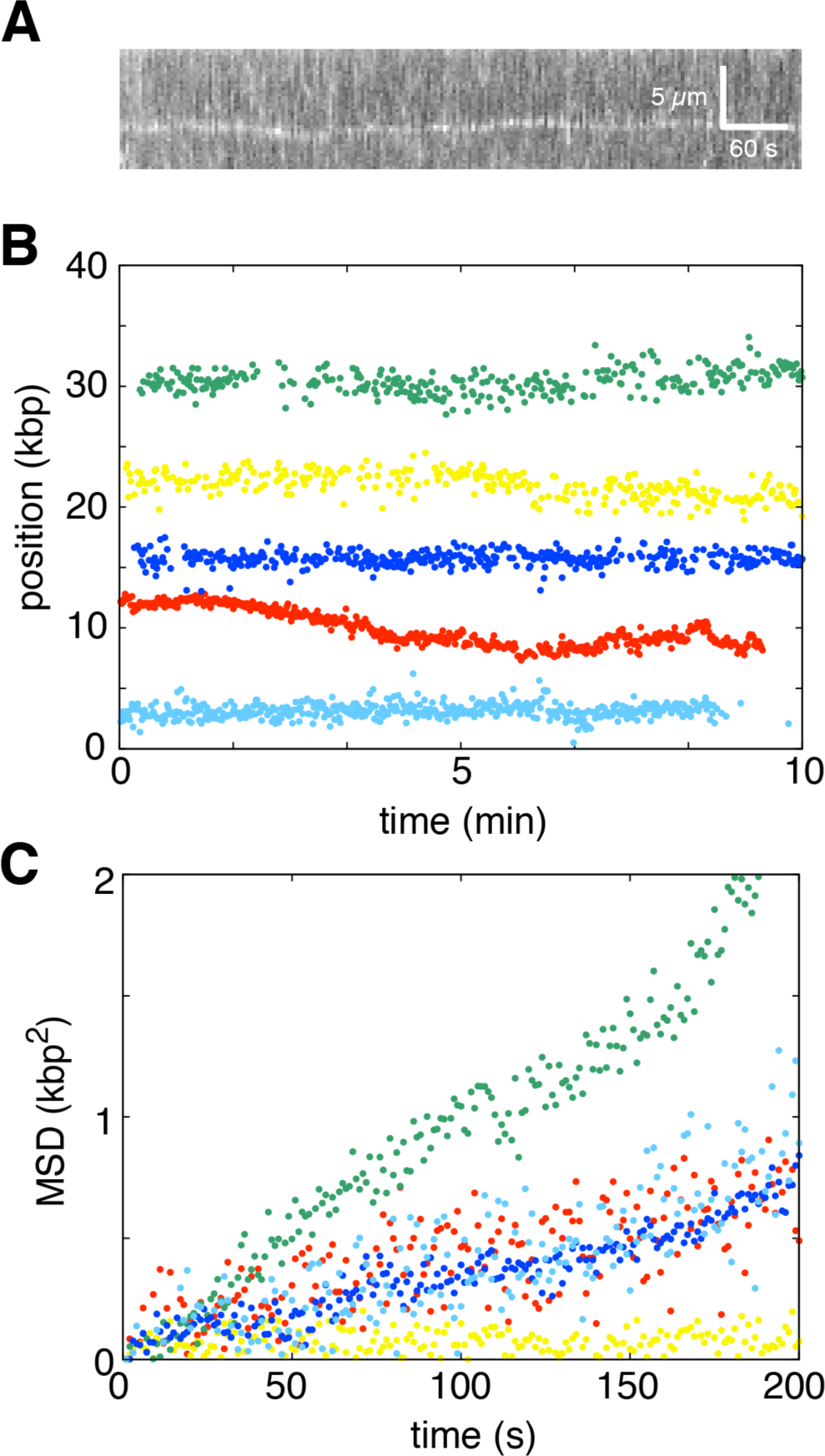
Condensin exhibits no motor activity in reactions with ATPγS. (**A**) Kymograph showing condensin bound to DNA in the presence of 4 mM ATPγS. (**B**) Examples of particle tracking data, and (**C**) MSD plots for data collected with wild-type condensin in reactions with 4 mM ATPγS.

**Fig. S5.**
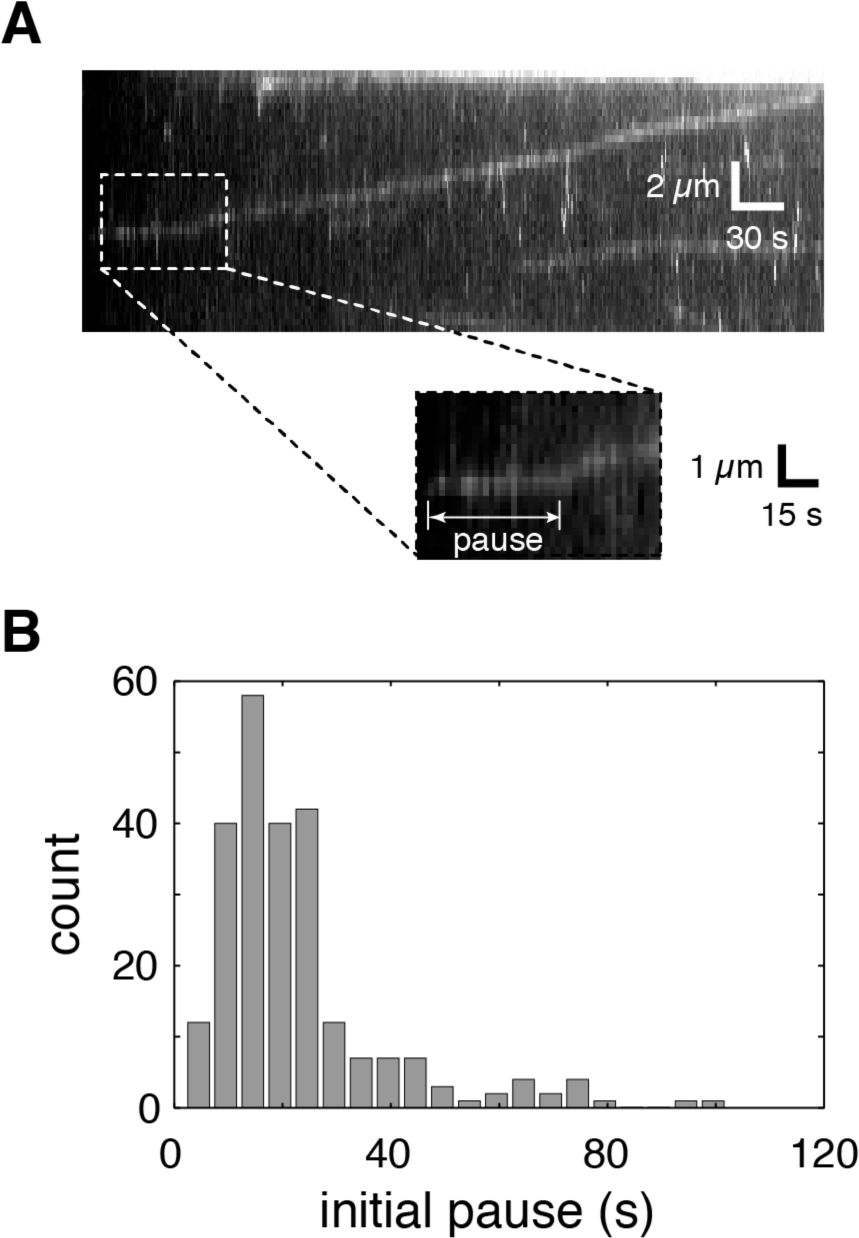
Condensin pauses prior to initiating translocation. (**A**) Kymograph highlighting the initial pause (τ_*pause*_) prior to the initiation of translocation (also see Fig. 2C). (**B**) Histogram showing the distribution of initial pause times prior to initiating translocation for reactions containing 4 mM ATP.

**Fig. S6.**
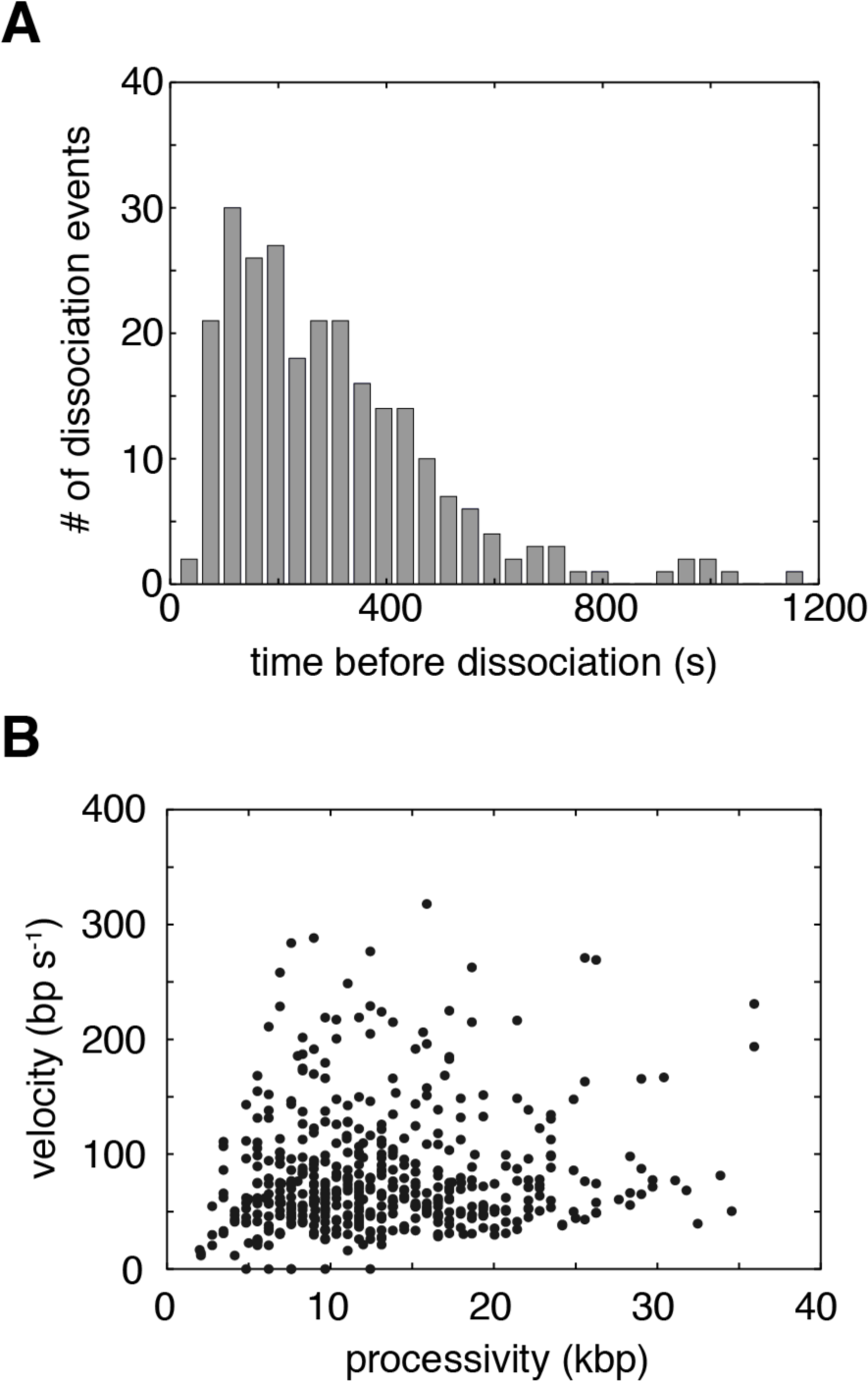
Condensin’s DNA-binding properties. (**A**) Distribution of binding lifetimes for translocating condensin complexes. (**B**) Scatter plot showing that there is no apparent correlation between condensin translocation velocity and processivity. All data shown in this figure reflect results from experiments conducted in the presence of 4 mM ATP.

**Fig. S7.**
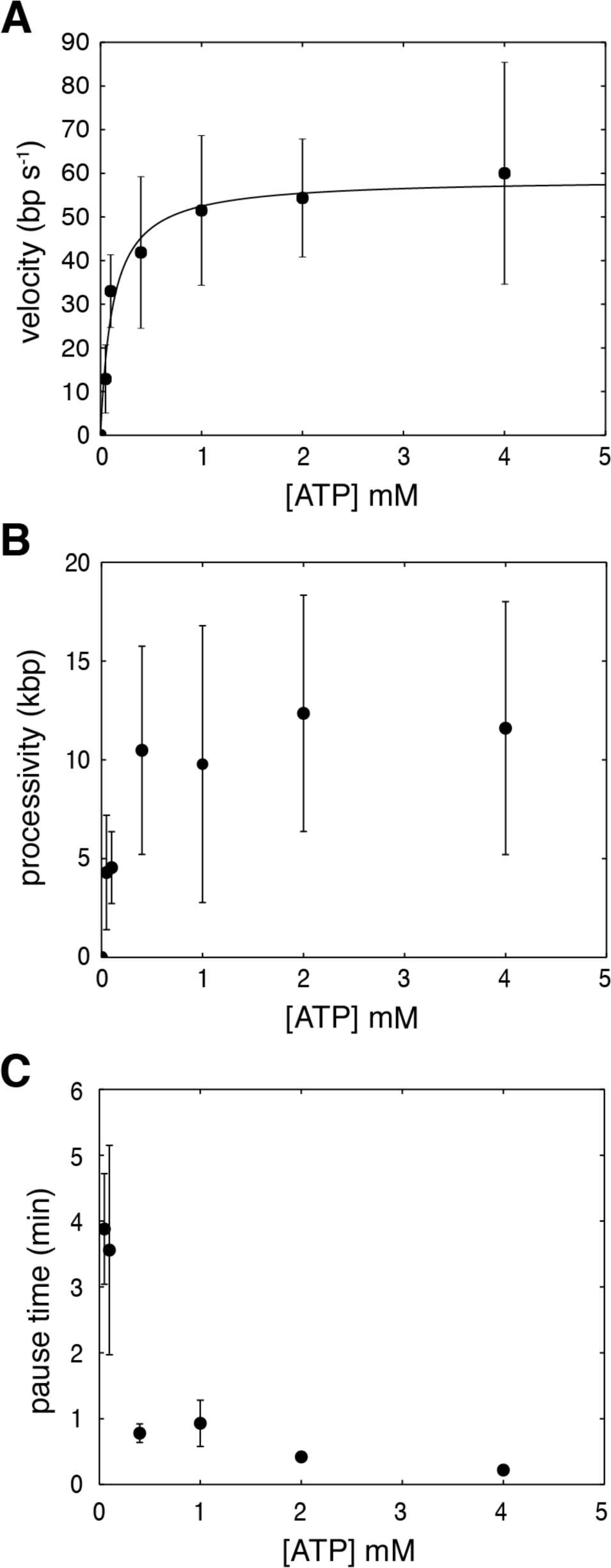
ATP concentration dependence of condensin translocation characteristics. (**A**) Condensin translocation velocity versus ATP concentration for data collected at room temperature (∼25°C). The data are fit to the Michaelis-Menton equation to extract the kinetic parameters *K*_*m*_ and *v*_*max*_. (**B**) Condensin processivity at different ATP concentrations, as indicated. (**C**) Initial condensin pause times (τ_*pause*_) prior to initiating translocation at different ATP concentrations. For each graph, error bars represent standard deviations calculated by boot strap analysis.

